# Improving water quality does not guarantee fish health: effects of ammonia pollution on the behaviour of wild-caught pre-exposed fish

**DOI:** 10.1101/2020.11.23.393868

**Authors:** Patricia Soler, Melissa Faria, Carlos Barata, Eduardo García-Galea, Beatriz Lorente, Dolors Vinyoles

## Abstract

Ammonia is a pollutant frequently found in aquatic ecosystems. In fish, ammonia can cause physical damage, alter its behaviour and even cause death. Exposure to ammonia also increases fish physiological stress, which can be measured through biomarkers. In this study, we analysed the effect of sublethal ammonia concentrations on the behaviour and the oxidative stress of *Barbus meridionalis* that had been pre-exposed to this compound in the wild. Wild-caught fish from a polluted site (pre-exposed fish) and from an unpolluted site (non-pre-exposed fish) were exposed, under experimental conditions, to total ammonia concentrations (TAN) of 0, 1, 5 and 8 mg/L. Swimming activity, feeding behaviour and oxidative stress response based on biomarkers were analysed. Pre-exposed fish showed both an altered behaviour and an altered oxidative stress response in the control treatment (0 mg/L). Differences in swimming activity were also found as pre-exposed fish swam less. Lower feeding activity (voracity and satiety) and altered response to oxidative stress were also observed at ≥ 1 mg/L TAN. Biomarker results confirmed pre-exposed fish suffer from a reduction in their antioxidant defences and, hence, showed increased oxidative tissue damage. In summary, pre-exposed fish showed more sensitivity to ammonia exposure than fish from a pristine site.

## Introduction

Ammonia is a commonly found pollutant in aquatic environments around the world [1, 2]. This compound can be found naturally, but there is also an additional contribution from sewage effluents, industrial waste and agricultural run-off [1]. The presence of ammonia in freshwater has been associated with the acidification of rivers and lakes, eutrophication and direct toxicity to aquatic organisms [1–3]. The toxicity of this compound on aquatic organisms will depend on the chemical form of ammonia, pH and temperature [4]. Furthermore, it will depend on the time of exposure [4]. This compound damages the gills, liver, kidney, spleen and other organ’s tissues of fish, therefore causing breathing difficulties [5, 6]. This may lead to physiological alterations and, eventually, exhaustion or death [6]. Ammonia can cause cell damage and can also affect the antioxidant defence system thus altering the levels of oxidative stress in fish [7, 8]. Ammonia can also alter fish behaviour. Fish exposure to sub-lethal concentrations of ammonia can reduce swimming activity [9], foraging behaviour [10] and the ability to flee from predators [10, 11].

Behavioural analyses are commonly used in ecotoxicology as indicators of sub-lethal toxicity in aquatic animals, and an increasing body of evidence has demonstrated the effectiveness of this approach in a wide range of exposure scenarios [12, 13]. Fish exposed to increased ammonia concentrations experience difficulty in eliminating this metabolite from the body [8] and, therefore, prolonged exposures to ammonia promotes its accumulation in fish [11]. Several studies indicated that fish pre-exposed to episodes of pollution by inorganic nitrogen compounds [14, 15] and heavy metals [16–18] could be more tolerant to these pollutants by acclimation. In these studies, it was shown that fish pre-exposed to sub-lethal concentrations of a pollutant exhibited an increased tolerance to exposure to high concentrations of the same pollutant. Fish pre-exposed to sub-lethal concentrations of ammonia pollution could tolerate high concentrations of this compound by increasing the ammonia excretion rate as well as by favouring the evolution of adaptive mechanisms [14, 15]. These mechanisms have also been shown to work with other types of stressors such as hypoxia [19], salinity [20] and temperature changes [21]. All these studies analyse the effect of fish pre-exposure from a biochemical and physiological point of view.

The aim of this study was to analyse, under experimental conditions, the effect of sublethal ammonia concentrations on the swimming activity and feeding behaviour of wild-caught fish that had been pre-exposed for a long time to this compound. The species selected for this study was the Mediterranean barbel, *Barbus meridionalis* (Risso 1827), a freshwater fish endemic to the NE Spain and SE France. Fish from a long-term polluted river and fish from a pristine stream were exposed to sublethal ammonia concentrations in the laboratory. Fish stress responses were complemented using biomarkers. The analysis of biomarkers may provide valuable information by assessing the activity of enzymes/markers involved in energy metabolism, detoxification, antioxidant defences and oxidative stress. In this study, biomarkers included lactate dehydrogenase (LDH), which is involved in anaerobic metabolism [22]; glutathione *S*-transferase, a xenobiotic that metabolizes II enzyme response [23]; glutathione (GSH) levels, which aid maintenance of the cell redox equilibrium as well as being a powerful antioxidant [24]; catalase (CAT EC 1.11.1.6—reduces H_2_O_2_ to water) an antioxidant enzyme involved in detoxifying reactive oxygen species and markers of oxidative tissue damage such as lipid peroxidation [25]. It has been suggested that *B. meridionalis* is relatively tolerant to organic pollution [26] and, globally speaking, more tolerant to pollution than other cyprinid species [26–28]. It was hypothesized that fish previously exposed to ammonia in the wild should have a higher tolerance to this compound than fish coming from unpolluted waters.

## Materials and Methods

### Study area and fish sampling

Two sites (polluted and unpolluted) were sampled in the Besòs River basin (NE Spain) (Fig 1). In both sites there was prior knowledge about the existence of a population of *B. meridionalis* [29, 30]. The polluted site was located in the Congost River, a 43 Km long tributary in the Besòs basin, 50 m downstream the Granollers WWTP (41°56’97.31”N, 2°27’15.66”E). The unpolluted site was located in the Castelló stream, a pristine 3 Km long tributary inside the San Llorenç del Munt i l'Obac Natural Park (41°65’16.97”N, 2°06’11.18”E). The concentration of total ammonia nitrogen (TAN) in the polluted site (Congost River) ranged from 0.54 mg/L to 24.70 mg/L between 2011 to 2015 (Table 1) (data provided for Granollers Town Council). In the unpolluted site (Castelló stream), the concentration of TAN ranged from 0.00 mg/L to 0.02 mg/L during the same period (Table 1) [31]. In this stream, there is no urban nucleus or any type of agricultural or industrial activity. Although ammonia is not the only pollutant present at these two sites, it is one of the most frequently found, not only in this river but in all rivers of NE Spain [2]. Table 1 shows the physical-chemical parameters analyzed at the two sites for the sampling month for Granollers Town Council (polluted site) and Fortuño et al. [32] (unpolluted site). Other contaminants such as contaminants of emerging concern (CEC) (pesticides, metals, industrial solvents, pharmaceuticals and personal care products), could be found in other sites across these basin [32].

**Table 1.** Physical-chemical water parameters from polluted and unpolluted sites.

|  |  | January 2016 |  | 2011 – 2015 |  |
| --- | --- | --- | --- | --- | --- |
| Physical-chemical parameters |  | Polluted | Unpolluted | Polluted | Unpolluted |
| General | pH | 8.0 | 8.0 | 8.0 ± 0.4 | 8.0 ± 0.3 |
|  | Oxygen (mg/L) | 7.8 | 5.5 | 8.1 ± 2.7 | 8.9 ± 2.4 |
| Salinity | Conductivity (µS/cm) | 1,112.0 | 444.7 | 1,062.5±107.5 | 613.9 ± 98.0 |
|  | Cl (mg/L) | 233.6 | 8.5 | 158.5 ± 40.8 | 13.2 ± 1.3 |
|  | SO<sub>4</sub> (mg/L) | 91.4 | 13.5 | 85.0 ± 7.8 | 16.7 ± 3.9 |
| Nutrients | NO<sub>2</sub> (mg/L) | 1.3 | 0.001 | 0.8 ± 0.7 | 0.007±0.003 |
|  | NO<sub>3</sub> (mg/L) | 39.5 | 0.01 | 26.0 ± 3.4 | 0.1 ± 0.1 |
|  | TAN (mg/L) | 4.8 | 0.02 | 2.4 ± 0.1 | 0.04 ± 0.02 |
|  | PO<sub>4</sub> (mg/L) | 5.3 | 0.003 | 4.1 ± 3.3 | 0.01 ± 0.01 |

**Fig 1.**
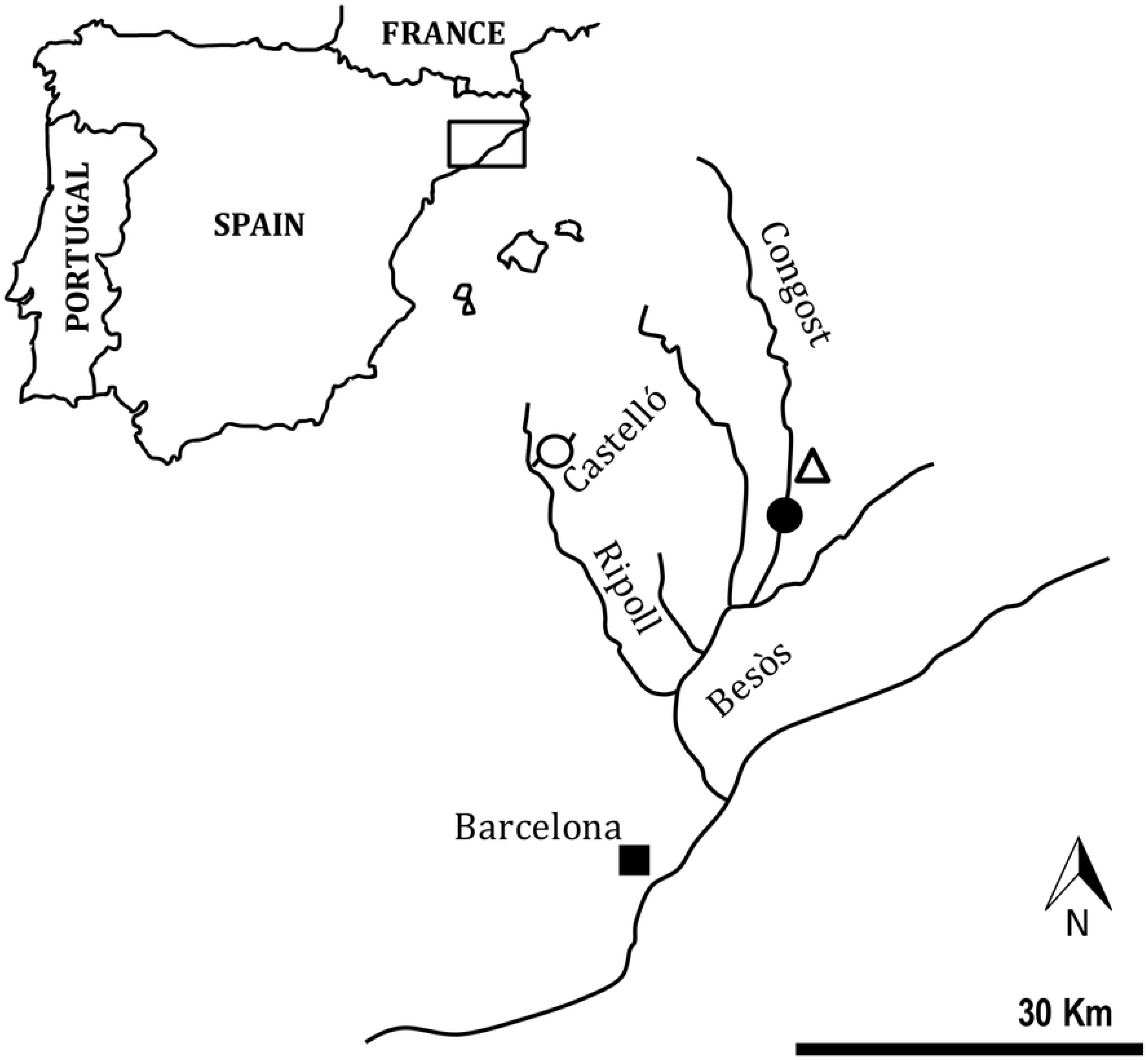
Map of the sampling sites in the Besòs River basin (NE of the Iberian Peninsula). The black point indicates the location of the polluted site in the Congost River (50 m downstream from a WWTP) where the pre-exposed fish were caught. The white point indicates the location of the unpolluted site in the Castelló stream.

Physical-chemical water parameters from both sites are shown for January 2016, when fish were sampled as well as the mean values for the period 2011 – 2015 (mean ± SD). All these data were provided by the Granollers Town Council (polluted site) and by Fortuño et al. (2018) (unpolluted site).

Fish were sampled by electrofishing using a portable unit which generated up to 200 V and 3 A pulsed D. C. A total of 72 individuals (40 in the polluted site and 32 in the unpolluted site) ranging from 5.5 to 10.8 cm were caught in January 2016. No differences in furcal length (FL, mean ± SD = 7.79 ± 1.31 cm) were found between the fish of the two sites. Once in the laboratory, fish of each site were acclimatized separately in 260 L aquaria over 21 days in clean dechlorinated water (there were 10-12 fish per aquaria). Chlorine elimination was achieved by storing water from the drinking supply net in 200 L containers during 48 h. According to Kroupova et al. [33] fish affected by nitrite poisoning that were placed in clean water for over six days, recovered the normal haematological parameters. Therefore, a period of 21 days seemed sufficient for the fish from the polluted site to recover normal physiological parameters. Aquaria were set in an acclimated room (20°C) under a 12 h light: 12 h dark photoperiod. All 260 L aquaria had the same equipment (biological filter and air diffusor), substrate (mix of sand, gravel, and coral with a proportion 2:2:1) and enough artificial refugees (PVC tubes and plastic plants) for reducing fish stress. Fish were fed “*ad libitum*” twice a day with frozen red chironomid larvae. A periodical cleaning of aquaria and partial water renovation (one-third of the volume) were carried out every 24 h. Physiochemical water conditions (mean ± SD) were controlled daily in the 260 L aquaria (water temperature = 21.97 ± 0.98 °C, pH = 8.30 ± 0.27, NO_3_^−^ = 5.63 ± 1.70 mg/L, NO_2_^−^ = 0.00 ± 0.00 mg/L, NH_4_^+^ = 0.00 ± 0.00 mg/L, and water hardness = 10.50 ± 4.36). These parameters did not show significant differences between aquaria during fish acclimatization.

### Experimental design

After the acclimatization period, fish pre-exposed to ammonia pollution in the wild (hereafter, pre-exposed fish) and fish from the unpolluted site (hereafter, non pre-exposed fish) were exposed to four TAN treatments (0, 1, 5 and 8 mg/L) as follows: each fish was placed in individual 20-L aquaria (40 cm large x 20 cm height x 25 cm deep) and transferred to the room where the experiment was carried out. The aquaria were divided into four groups and a treatment was randomly assigned to each group (Fig 2). For the Congost river (pre-exposed fish) there were ten aquariums per treatment (n = 40), while for the Castelló stream (non pre-exposed fish), there were eight aquariums per treatment (n = 32). Aquaria were positioned in two rows, side by side, within each group (Fig 2). In order to reduce fish stress, the lateral walls between neighboring aquaria were left transparent. To avoid fish interaction with the environment, the external and frontal walls as well as the bottom of aquaria were covered by blue acetate sheets. Before starting the experiment, each fish was acclimatized to its 20 L aquaria for four days and fed daily with red chironomid larvae. During these four days of fish acclimatization, partial water changes were carried out every day, and TAN concentrations were measured with indophenol blue spectrophotometric method (in all aquaria TAN concentration was maintained at 0 mg/L; mean ± SD = 0.00 ± 0.00 mg/L).

**Fig 2.**
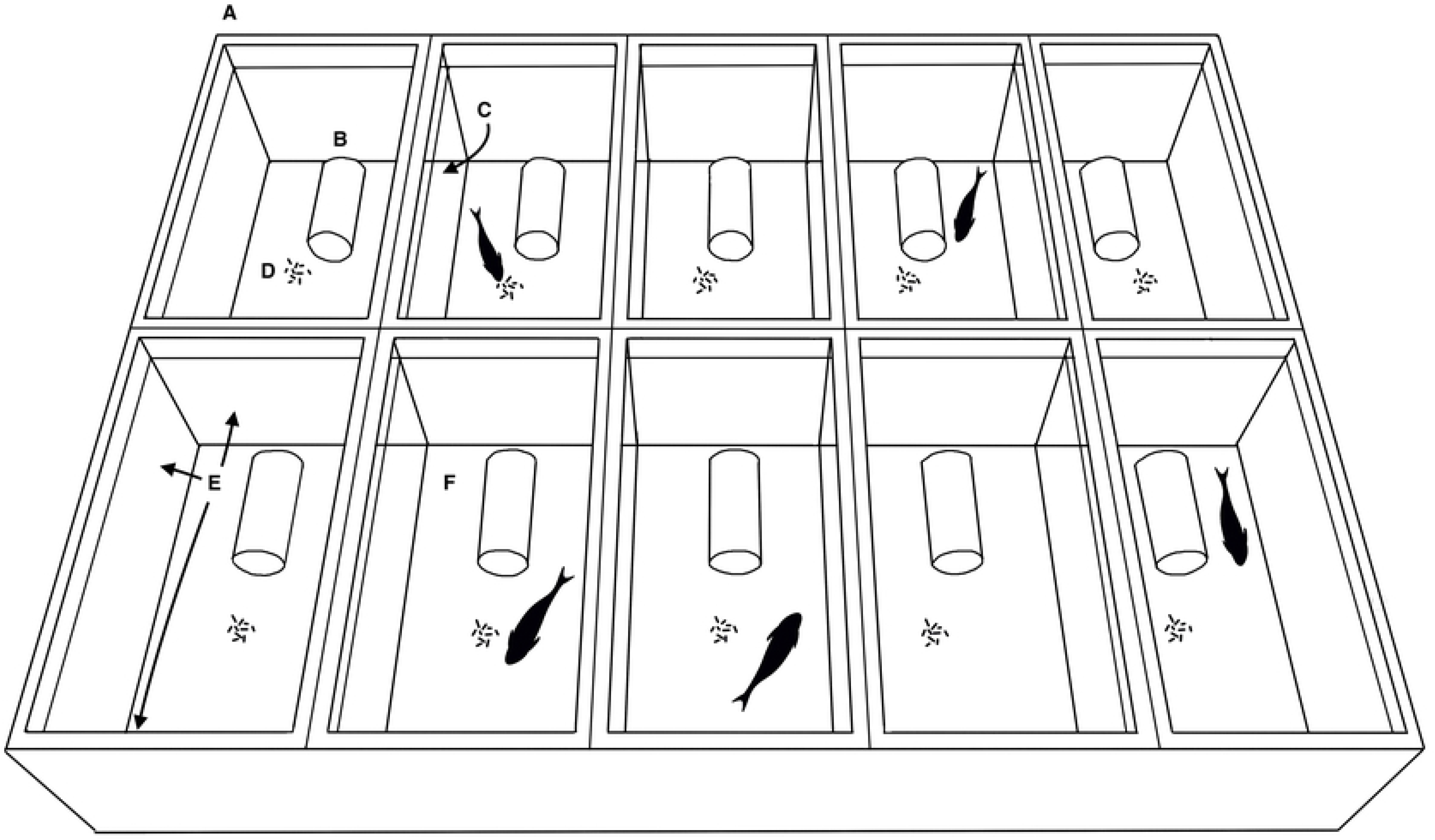
Individual experimental aquaria used for each TAN treatment. A: Schematic representation of a group of 20 L individual aquaria used for each TAN treatment in the experiment with *B. meridionalis* (aquaria were placed on a solid deck pallet to facilitate handling). B: PVC tube refuge. C: Transparent lateral walls between neighbouring aquaria. D: Red chironomid larvae used to observe feeding behaviour. E: Exterior and frontal walls of the aquaria covered with blue acetate sheets. F: Bottom of aquaria covered by a blue acetate sheet.

Next, fish were exposed to the assigned TAN treatment for eight days. The experiment was first carried out with the pre-exposed fish. After eight days, the experiment was repeated with the non pre-exposed fish. The TAN concentrations per experimental aquaria were achieved by adding analytical grade ammonium bicarbonate solutions (NH_4_HCO_3_, Sigma-Aldrich, Barcelona, Spain). These solutions were dispensed with automatic pipettes after water changes. A daily cleaning of aquaria and a two-third of the water volume renovation were carried out with dechlorinated water to guarantee the experimental conditions. TAN concentrations were measured daily by the indophenol blue spectrophotometric method. Once the absorbance values had been recorded for each sample, NH_4_^+^ concentration was calculated using the equation of the calibration curve and the proportion of the NH_3_ form was calculated following Thurston et al. [34] procedures. During the experiment, the aquaria group of each TAN treatment was visually isolated from the researchers with opaque curtains. In order to observe the activity of fish, a PVC tube (4 cm diameter x 13 cm length) was placed in each 20 L aquaria as a fish refuge (Fig 2). In order to observe feeding activity, fish were fed above satiation requirements (20 red chironomid larvae per fish were sufficient to quantify satiety). Fish behaviour was recorded with an overhead shot for each group of aquaria (TAN treatment) using a Sony HD (HDR-SR1E) camera. The experiment lasted for eight days and recordings were made on alternative days (four days) between 9:00 and 12:00 AM. Every day, the recording order of each group of aquaria (TAN treatment) was established at random. Fish were only fed during the recording days. Two behavioural variables per individual were analyzed from video recordings: swimming activity (during 10’) and feeding behaviour (until fish stopped eating). The swimming activity was analysed by three variables: (1) “Swimming”, amount of time during which fish make displacements of the body using body or fin movement as propulsion (s), (2) “Not visible”, amount of time during which the fish was not visible because it was remaining inside the shelter (s), and (3) “Resting”, amount of time fish spent lying motionless on the bottom of the aquaria (s). Total swimming activity was expressed as a percentage of the total observation time [35]. Feeding behaviour was analysed by measuring: (1) “Latency”, defined as the amount of time the fish took to start touching the food (s); (2) “Voracity”, defined as the number of chironomid larvae the fish ate in one minute and (3) “Satiety”, defined as the amount of time until the fish either stopped eating or they started spitting out the food (s).

The concentration of NH_3_ (mean ± SD, mg/L) for each TAN treatment was not significantly different between pre-exposed and non pre-exposed fish (GLM): [0 mg/L] = 0.007 ± 0.010, [1 mg/L] = 0.139 ± 0.077, [5 mg/L] = 0.534 ± 0.218, [8 mg/L] = 0.645 ± 0.237). Physiochemical parameters were controlled daily for each 20 L aquaria during the experiment. No differences in physiochemical parameters (mean ± SD) were found during the experiment between fish from the two sites and between the aquaria of each TAN group (GLM) (water temperature = 21.27 ± 0.45 °C, pH = 8.33 ± 0.18, NO_3_^−^ = 4.81 ± 0.68 mg/L, NO_2_^−^ = 0.00 ± 0.00 mg/L, and water hardness = 15.05 ± 3.76).

All applicable international, national, and/or institutional guidelines for the care and use of animals were followed. The scientific procedure of this work was approved by the Animal Ethic Committee of the University of Barcelona (registration N° 9296), which follows European Directive 2010/63/UE on the protection of animals used for scientific purposes. One of the co-authors holds a category C FELASA certificate that regulates the use of animals for experimental and other scientific purposes.

### Biochemical determination

For the biochemical determinations, fish were anesthetized on ice at the end of the experiment and euthanatized by decapitation. Biomarkers were analysed in the liver tissue for each individual fish according to Faria et al. [36].

The following reagents were purchased from Sigma-Aldrich (St. Louis, MO, USA): potassium phosphate dibasic (K_2_HPO_4_); potassium phosphate monobasic (KH_2_PO_4_); potassium chloride (KCl); ethylenediamine-tetraacetic acid, disodium, salt, dihydrate (EDTA); hydrogen peroxide (H_2_O_2_); reduced glutathione (GSH); sodium azide; 1-chloro–2,4–dinitrobenzene (CDNB); glutathione S-transferase, from equine liver (GST) (EC 2.5.1.18); monochlorobimane (mCB); sodium pyruvate; β-Nicotinamide adenine dinucleotide, reduced dipotassium salt (NADH), 2,6-di-tert-butyl-4-methylphenol (BHT); 1-methyl-2-phenylindole (MPI); 1,1,3,3-tetramethoxypropane (TMP) and Bradford reagent. All the other chemicals were analytical grade and were obtained from Merck (Darmstadt, Germany).

Except for catalase activity, where a cuvette assay was used (Life Science UV/Vis Spectrophotometer DU® 730, Beckman Coulter – Fullerton, CA, USA), all the bioassays were performed in microplates (Synergy 2 Multi-Mode Microplate Reader, BioTek® Instuments – Vermont, USA).

Liver tissue was homogenized in ice-cold 0.1M phosphate buffer with 150mM KCl and 0.1mM ethylenediamine-tetraacetic acid, disodium, salt, dihydrate (EDTA), then centrifuged at 10 000xg, 4°C for 30 minutes. The supernatant was collected, aliquoted and stored at −80°C for biomarker determination.

CAT activity was measured by estimating the decrease in absorbance at 240 nm due to H_2_O_2_ (50 mM H_2_O_2_ in 80 mM phosphate buffer, pH 6.5) consumption (extinction coefficient 40 M^−1^cm^−1^) according to Aebi [37]. Reaction volume and time were 1 mL and 1min, respectively. GST activity towards CDNB was measured as described by Habig et al. [38]. The reaction mixture contained 0.1M phosphate buffer (pH 7.4), 1 mM CDNB and 1 mM GSH. The formation of S – 2,4 dinitro phenyl glutathione conjugate was evaluated by monitoring the increase in absorbance at 340 nm during 5 minutes. Enzyme activity was determined using GST’s extinction factor coefficient of 9.6 mM^−1^ cm^−1^. Results were normalized by tissue total assay protein content. Reduced glutathione (GSH) quantification was adapted from zebra mussel digestive gland according to Kamencic et al. [39]. It consists on adding 0.1mM of mCB along with 1U/ml of GST to each sample. Then the GSH present in the cells forming a GSH-mCB complex is measured fluorometrically at excitation: emission wave length of 360:460 nm, after an incubation period of 90 minutes at room temperature and protected from light. The total content of GSH was then extrapolated from a GSH standard curve determined under the same physical and chemical conditions as the samples, and the results were normalized by the tissue wet weight (g ww).

Lactate dehydrogenase (LDH) activity was determined according to Diamantino et al. [22] by monitoring the absorbance decrease at 340 nm due to NADH oxidation. The reaction contained 100 mM phosphate buffer (pH 7.4), 0.15 mM NaOH, 1.18 mM pyruvate and 0.18 mM NADH.

Lipid peroxidation (LPO) was determined by quantifying the levels of malondialdehyde (MDA) according to Esterbauer et al. [40]. The MDA assay was based on the reaction of the chromogenic reagent 1-methyl-2-phenylindole with MDA at 45°C, giving rise to a chromophore with absorbance at 586nm. Samples were incubated with 5mM 1-methyl-2-phenylindole in acetonitrile:methanol (3:1 v/v), 5.55% of HCl and 0.01% BHT at 45°C, for 40 minutes. Absorbance was read at 560nm and MDA content in each sample was extrapolated from the standard curve of 1,1,3,3-tetramethoxypropane (TMP) treated under similar conditions as samples. The final results were normalized by tissue wet weight (g ww).

Total protein concentrations were accessed by the Bradford method using bovine serum albumin (BSA) as a standard [41].

### Statistical analyses

Differences between TAN treatments and sites were analysed by means of a generalized lineal mixed model (GLMM). For swimming activity, “Swimming” and “Not visible” were analysed separately and used as dependent variable. The variable “Resting” was not analysed as it was a complementary variable to the other two. For feeding behaviour, “Latency”, “Voracity” and “Satiety” were analysed separately and used as dependent variables. In all cases “site” (2 levels: pre-exposed and non pre-exposed fish) and “TAN treatments” (4 levels: “0”, “1”, “5” and “8” mg/L TAN) were used as factors together with their interaction. The gamma distribution was assumed in the analysis of swimming activity and the Poisson distribution was assumed in the analysis of feeding behaviour. The variable “Individual” was added to the model as a random factor.

Biomarker responses across fish from the two sites (pre-exposed and non pre-exposed fish) and TAN treatments were analysed through a lineal model (LM) with the same factors (“site” and “TAN treatments”) [42]. Differences between TAN treatments against control ones were further compared using Dunnett’s post-hoc test [42].

All analyses were conducted with R 3.4.3 [43]. GLMM assuming a Poisson or a gamma distribution was performed using glmer() (package “lme4”: [44]). Non-significant interactions were removed from final models. Homogeneity and normality of residuals were visually checked for all models. All significant differences are *P* ≤ 0.05.

## Results

### Behavioural variables

The “Swimming” and “Not visible” GLMM models showed no significant effect of TAN treatments within and across sites (interaction) (*P* > 0.05). Only a significant effect of site (pre-exposed and non pre-exposed fish) (*P* < 0.001) was shown for these two variables. The swimming activity of fish that had been pre-exposed to ammonia pollution in the field was lower than that of non pre-exposed fish. Non pre-exposed fish swam for a longer time (67% of the time; mean = 643.89 s; 95% confidence interval = 504.10 – 891.34) than pre-exposed fish (57.3% of the time; mean = 548.92 s; 95% confidence interval = 444.92 – 716.43) regardless of the TAN treatments they were in. Similarly, non pre-exposed fish spent significantly less time hidden inside the shelter (mean = 206.76 s; 95% confidence interval = 150.01 – 332.90) than pre-exposed fish (mean = 232.99 s; 95% confidence interval = 165.76 – 392.43).

The analysis of feeding behaviour showed no significant effects of the interaction between TAN treatment and site for none of the variables (*P* > 0.05). For “Latency” GLMM model showed a significant effect between sites (*P* < 0.002) but not for TAN treatments (*P* > 0.05). Non pre-exposed fish had a higher latency than pre-exposed fish (Fig 3A). In contrast, GLMM models for “Voracity” and “Satiety” variables showed a significant effect between sites and TAN treatments within each site (*P* < 0.001). Non pre-exposed fish had a higher voracity (Fig 3B) and were satiated later (Fig 3C) than pre-exposed ones. In both cases (sites), significant differences were found between the control TAN concentration (0 mg/L) and the three TAN treatments (1, 5, 8 mg/L) for “Voracity” and “Satiety” variables.

**Fig 3.**
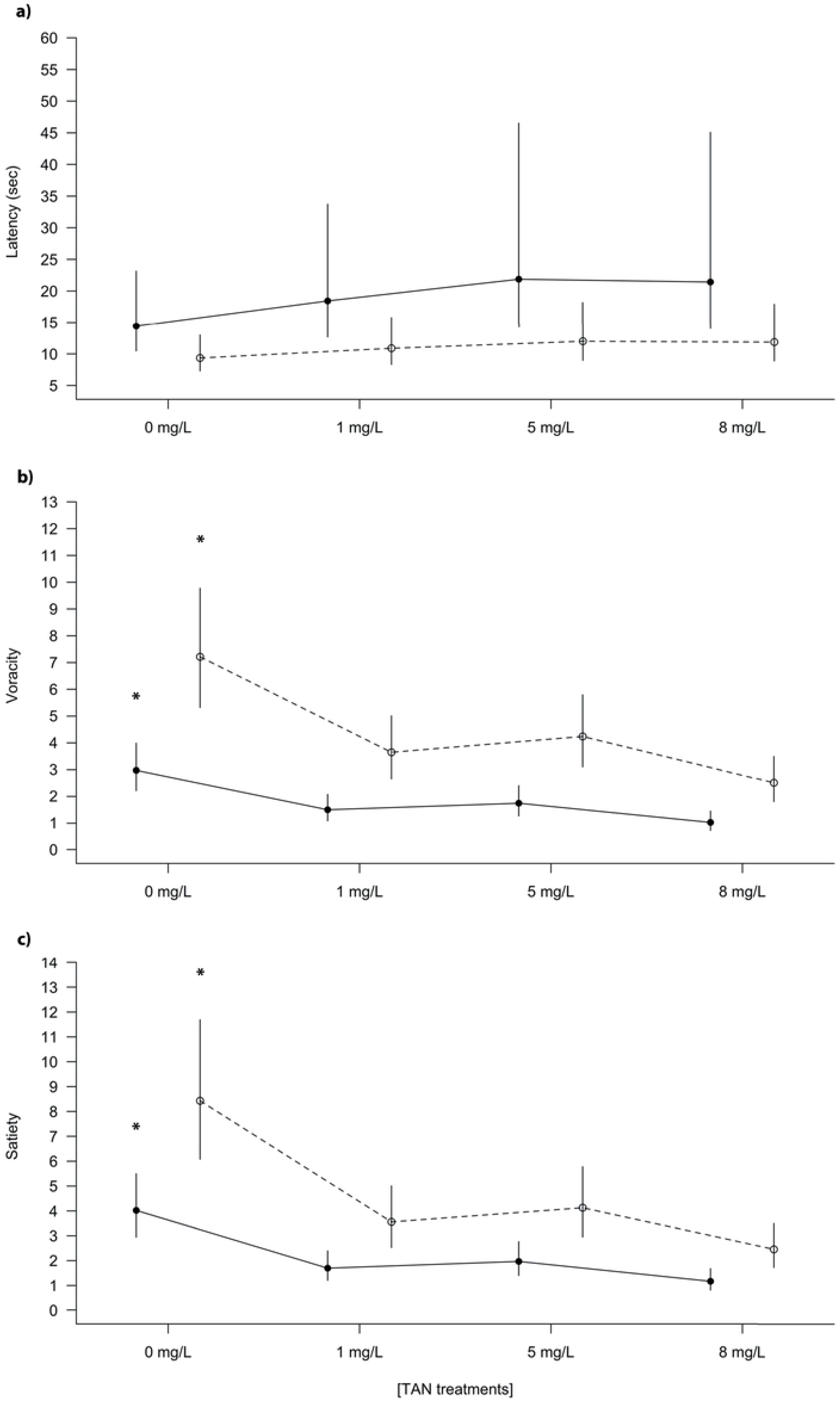
Feeding behaviour of *B. meridionalis* exposed to the TAN treatments during the experiment. (a) Latency (time taken to start touching the food), (b) Voracity (number of red chironomid larvae eaten in one minute) and (c) Satiety (total number of red chironomid larvae fish eaten) are shown for the pre-exposed fish from the polluted site (black points and solid lines), and the non pre-exposed fish from the unpolluted site (white points and dashed lines). * indicates significant differences (*P* < 0.05) between the control concentration and the TAN treatments, following a generalized linear mixed model (GLMM).

### Biochemical determination

The results of the analysis of biomarkers show that there were significant (*P* < 0.05) differences between sites (pre-exposed and non pre-exposed fish) in three out of the five studied biomarkers (Table 2, Fig 4). TAN treatment within and across sites (interaction) also affected the activities of CAT, GST and levels of LPO. Pre-exposed fish had lower CAT activities and lower levels of GSH, and the activities of CAT and levels of LPO increased across TAN treatments. In fish from both sites the activities of GST were enhanced at 1 mg/L of TAN (Fig 4).

**Table 2.**
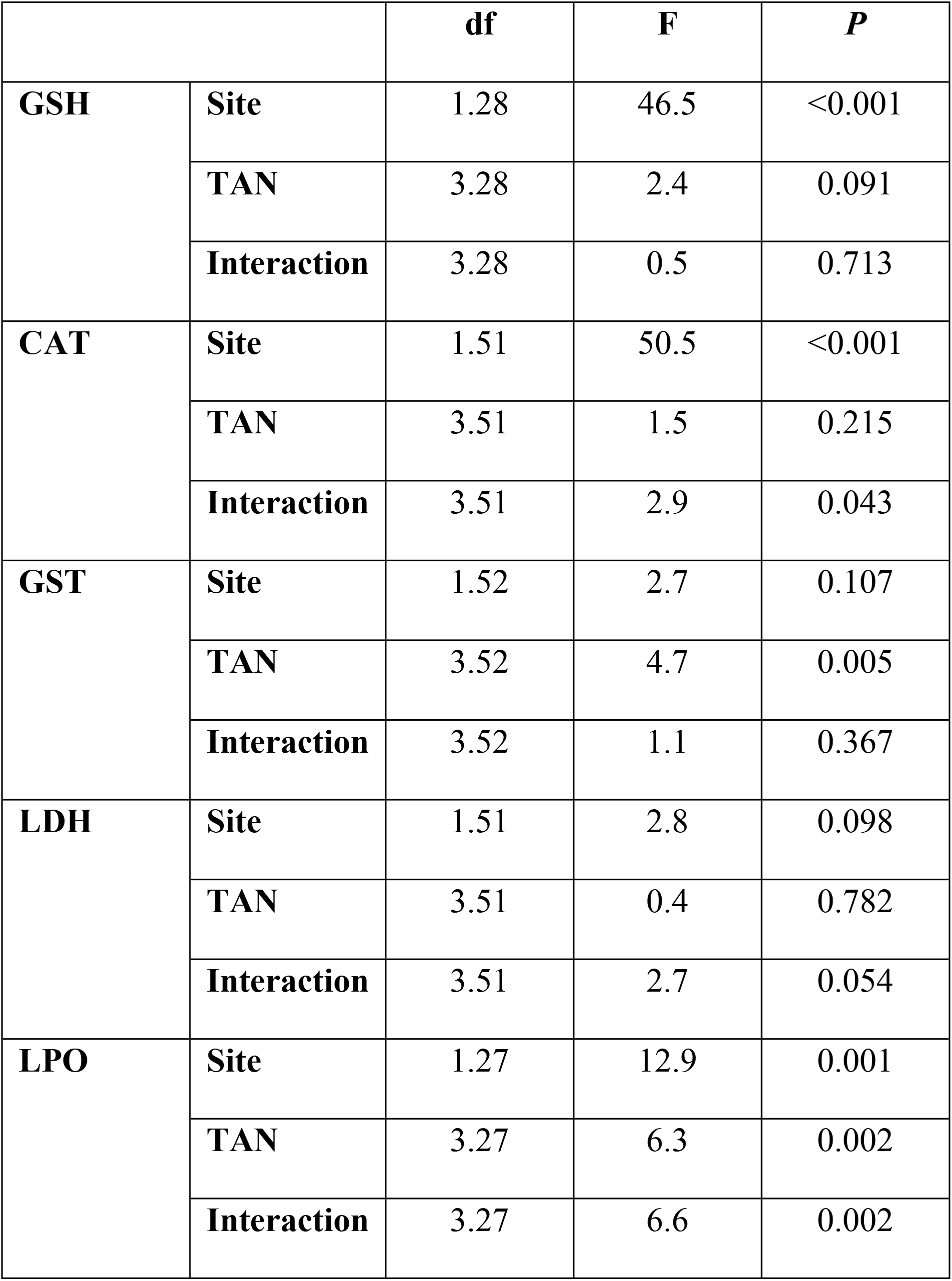
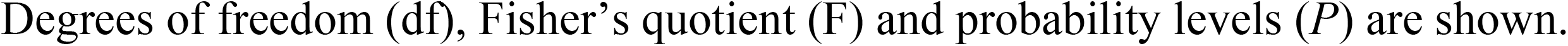
Linear model results for testing the effects of site and TAN on the studied biomarkers in *B. meridionalis*.

|  |  | df | F | P |
| --- | --- | --- | --- | --- |
| **GSH** | **Site** | 1.28 | 46.5 | <0.001 |
|  | **TAN** | 3.28 | 2.4 | 0.091 |
|  | **Interaction** | 3.28 | 0.5 | 0.713 |
| **CAT** | **Site** | 1.51 | 50.5 | <0.001 |
|  | **TAN** | 3.51 | 1.5 | 0.215 |
|  | **Interaction** | 3.51 | 2.9 | 0.043 |
| **GST** | **Site** | 1.52 | 2.7 | 0.107 |
|  | **TAN** | 3.52 | 4.7 | 0.005 |
|  | **Interaction** | 3.52 | 1.1 | 0.367 |
| **LDH** | **Site** | 1.51 | 2.8 | 0.098 |
|  | **TAN** | 3.51 | 0.4 | 0.782 |
|  | **Interaction** | 3.51 | 2.7 | 0.054 |
| **LPO** | **Site** | 1.27 | 12.9 | 0.001 |
|  | **TAN** | 3.27 | 6.3 | 0.002 |
|  | **Interaction** | 3.27 | 6.6 | 0.002 |

Degrees of freedom (df), Fisher's quotient (F) and probability levels (*P*) are shown.

**Fig 4.**
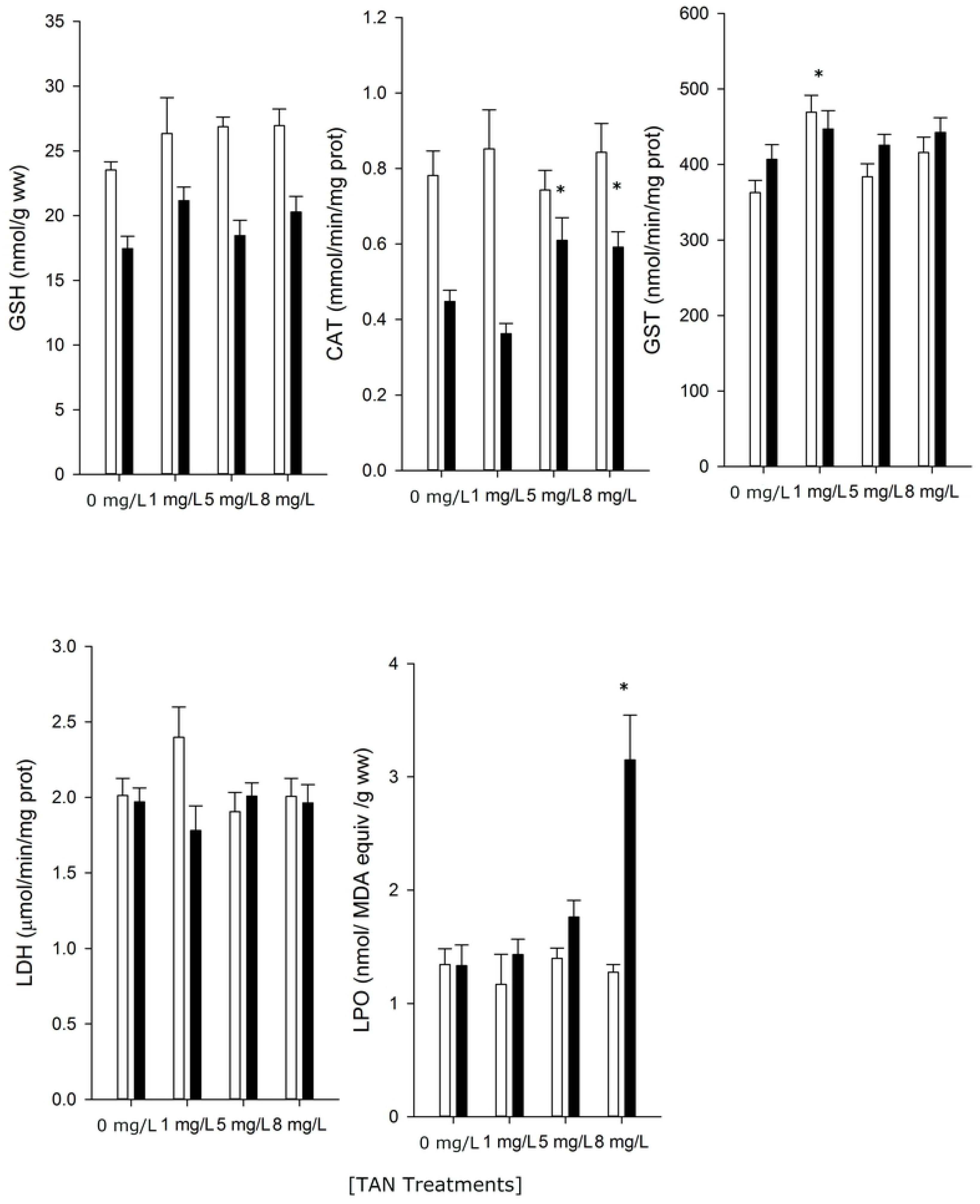
Antioxidant and oxidative stress responses (Mean ± SE, N=10) of *B. meridionalis* exposed to increasing TAN concentrations during the experiment. Black and white bars indicate the results for the fish from the polluted site (pre-exposed fish) and the unpolluted site (non pre-exposed fish), respectively. * indicates significant differences (*P* < 0.05) within each site from control TAN treatment (0 mg/L TAN) following LM model and Dunnett’s post-hoc tests.

## Discussion

Chronic exposure to pollution by nitrogen inorganic compounds (NH_4_^+^, NH_3_, NO_2_^−^, HNO_2_ and NO_3_^−^) has effects on the reproduction, growth and survival of freshwater fish [2]. Specifically, exposure to NH_4_^+^ and NH_3_ (TAN) pollution can cause gill damage, anoxia, disruption of blood vessels and osmoregulatory activity (damage to the liver and kidneys), and a decrease in the effectiveness of the immune system [1]. In addition, NH_4_^+^ ions contribute to an internal reduction of Na^+^ which, in turn, increases the toxicity by NH_3_ [2]. All these effects can result in a reduction in fish feeding activity, fecundity and survival, leading to a reduction of the size of populations [2].

In the present study, wild-caught fish pre-exposed for a long-term period to ammonia pollution in a contaminated river near a WWTP showed an altered behaviour and suffered from an increased physiological stress as compared to non pre-exposed fish from a pristine stream. Analysis of fish swimming activity showed that, regardless of the TAN treatments, pre-exposed fish were less active and spent more time hiding in the refuge than non pre-exposed fish. The only studies on the effects of ammonia on fish swimming activity have been conducted on salmonids, in laboratories or farms. According to Tudorache et al. [10] and Wicks et al. [9], the swimming activity of salmonids is reduced at concentrations between 0.2-1 mg/L of TAN (that is, at 0.009 – 0.04 mg/L NH_3_). Pre-exposed fish spending more time inside the PVC shelters (“not visible” time) might indicate that these fish had their exploratory activity altered. A decrease in the exploratory activity has been reported in several fish species exposed to crude-oil pollution [45], pesticides [46] and pharmaceutical products [47].

The feeding behaviour of *B. meridionalis* was also altered. Pre-exposed fish had lower voracity than non pre-exposed fish regardless of the TAN treatments (0, 1, 5 and 8 mg/L TAN). Within each site (pre-exposed and non pre-exposed fish), lower voracity was observed from the lowest TAN concentration (1 mg/L). A reduction in voracity has been reported for salmonids under TAN concentrations from 1 to 3 mg/L [10, 48, 49]. According to Schram et al. [6], in a non salmonid fish (*Clarias gariepinus*), food consumption was also drastically reduced at TAN concentrations higher than 1 mg/L. In the present study, latency (the time that the fish took to start touching the food) was lower in pre-exposed fish, regardless of the TAN treatment. A low latency has been related to a low capacity to find food and capture prey [50]. Furthermore, several studies relate lower latency with lower efficiency to flee from predators [11, 51, 52].

The present study was conducted under a concentration of ammonia within the range of LC_50_ (the tested range was from 0.007 to 0.645 mg/L NH_3_). However, the tolerance to NH3 in cyprinids could be higher. The LC_50_ for cyprinids ranked between 0.685 and 1.720 mg/L NH_3_ [1]. For cyprinids, the sublethal concentrations in which negative physiological effects begin to be observed has been described in a range of 0.105-0.247 mg/L NH_3_ [53–55]. The limits of tolerance to this and other compounds are variable depending on each fish species so that it would be necessary to investigate their effects under natural conditions.

Antioxidant enzyme activities such as those of CAT and reduced glutathione has been reported to be important antioxidant mechanisms against oxidative stress-mediated effects of ammonia in fish [8, 56–66]. Pre-exposed fish (from the polluted site) had lower constitutive levels of the above mentioned antioxidant defences and consequently were unable to detoxify the excess of reactive oxygen species (ROS) generated by ammonia, leading to enhanced tissue levels of oxidative damage measured as LPO. Interestingly, only in fish from the polluted site (pre-exposed fish), the activities of CAT increased in individuals, exposed to ammonia, thus indicating that the exposure to this compound increased ROS and, hence, triggered the antioxidant defences of these fish. In fish from the unpolluted site (non pre-exposed fish), the high constitutive levels of antioxidant defences protected them from ROS generated by ammonia. [8] Sinha et al. (2014) reported that fish species intolerant to ammonia, such as trout, rely mainly on glutathione-dependent defensive mechanisms, while more tolerant species, such as carps, utilize antioxidant enzymes such as CAT and ascorbate. High tolerant species, such as goldfish, use both of these protective systems, and show more effective anti-oxidative compensatory responses towards oxidative stress induced by ammonia [8]. Thus, our results are in line with previous studies, as *B. meridionalis* considered a tolerant species to ammonia.

Results in this study indicated that the exposition of fish to high ammonia concentrations did not guarantee, at least short term, the recovery of a good health status and/or a greater tolerance to a high concentration of this compound. Fish pre-exposed to ammonia pollution in the wild showed an altered behaviour at the control concentration (0 mg/L TAN). This could be a consequence of pre-existing physiological problems due to exposure to ammonia and other pollutants in nature [67–69]. The feeding behaviour and the response to the oxidative stress of *B. meridionalis* (both pre-exposed and no pre-exposed fish) follow the same pattern, reacting equally to the first 1 mg/L TAN treatment. However, pre-exposed fish had a more marked response in feeding behaviour and biomarkers under the different treatments of TAN. A reduction in food intake is directly related to both a lower growth and a low rate of protein synthesis [70]. Reported studies have shown that low protein synthesis rates represent a large proportion of energy costs in fish, and this has a direct impact on the growth efficiency of individuals [71]. Alteration of the behaviour parameters analysed in this study can be extrapolated to other traits such as exploration activity, boldness and ability to avoid predators [72]. Ammonia can affect social interactions as well, by altering dominance relationships, hierarchical dynamics and predator-prey relationships [10, 73].

Ammonia pollution is a common problem in freshwater ecosystems [7]. Despite the efforts of implementing the [74] European Water Framework Directive (2000/60/RC) (2000) there are still many WWTPs that do not have tertiary purification systems of urban wastewater, which leads to an increase in nitrogen compounds in aquatic ecosystems [75, 76]. Improving water quality is an important key to enhance the conservation of river ecosystems. However, our results indicated that fish that previously survived in a polluted environment did not recover their health in more purified waters. In summary, although habitats are improving their environmental quality, the survival of fish populations that have been pre-exposed to contamination could be compromised. In freshwater ecosystems, which have suffered an 83% decline in vertebrate populations from 1970 to 2014 [77], all factors affecting the survival of individuals are of great relevance.

## Conclusions

*B. meridionalis* pre-exposed in the wild to pollution by ammonia presented a swimming activity, a feeding activity and the response to oxidative stress altered when placed in non contaminated water under experimental conditions. Feeding behaviour and the biomarker response of *B. meridionalis* was affected by ammonia and pollution history. Pre-exposed fish (from a polluted site) had less voracity and satiated before fish from an unpolluted site (non pre-exposed fish). In addition, pre-exposed fish were more affected by the different TAN treatments and these alterations appeared from the lowest concentration of TAN (1 mg/L). The results of swimming activity showed that pre-exposed fish spent less time swimming and more time hidden. However, this behavioural response was not related to the different TAN treatments and could be related to the damage caused by pre-exposure to ammonia and other pollutants present in the river. Fish pre-exposed to ammonia in the wild also had the antioxidant defences depressed and consequently were less tolerant to high concentrations of ammonia. Therefore, our results indicated that the recovery of water quality is not necessarily related to the restoration of fish health. There is a physiological cost of being adapt to pollution present in rivers.

## Acknowledgements

We thank I. Ramirez and F. López of the Department of Biochemistry and Molecular Biomedicine (University of Barcelona), for giving us access to their laboratory. We thank P. Fortuño (University of Barcelona) and X. Romero (biologist and superior technician of environment and natural environment of the Granollers Town Council) for provided data. We also thank J. Guinea and P. Manning for the assistance in the laboratory tasks. The authors are grateful to V. Bonet for the English review. This research did not receive any specific grant from funding agencies in the public, commercial, or not-for-profit sectors.

